# CNEr: a toolkit for exploring extreme noncoding conservation

**DOI:** 10.1101/575704

**Authors:** Ge Tan, Dimitris Polychronopoulos, Boris Lenhard

## Abstract

Conserved Noncoding Elements (CNEs) are elements exhibiting extreme noncoding conservation in Metazoan genomes. They cluster around developmental genes and act as long-range enhancers, yet nothing that we know about their function explains the observed conservation levels. Clusters of CNEs coincide with topologically associating domains (TADs), indicating ancient origins and stability of TAD locations. This has suggested further hypotheses about the still elusive origin of CNEs, and has provided a comparative genomics-based method of estimating the position of TADs around developmentally regulated genes in genomes where chromatin conformation capture data is missing. To enable researchers in gene regulation and chromatin biology to start deciphering this phenomenon, we developed *CNEr*, a R/Bioconductor toolkit for large-scale identification of CNEs and for studying their genomic properties. We apply *CNEr* to two novel genome comparisons - fruit fly vs tsetse fly, and two sea urchin genomes - and report novel insights gained from their analysis. We also show how to reveal interesting characteristics of CNEs by coupling CNEr with existing Bioconductor packages. *CNEr* is available at Bioconductor (https://bioconductor.org/packages/CNEr/) and maintained at github (https://github.com/ge11232002/CNEr).

## Introduction

Conserved Noncoding Elements (CNEs) are a pervasive class of extremely conserved elements that cluster around genes with roles in development and differentiation in Metazoa [1,2]. While many have been shown to act as long-range developmental enhancers [3,4], the source of their extreme conservation remains unexplained [5,6]. The need to maintain arrays of CNEs in cis to the genes they regulate has led to their spatial arrangement into clusters termed Genomic Regulatory Blocks (GRBs) [7,8]. The role of those clusters in genome organisation is suggested by recent findings demonstrating that ancient metazoan clusters of extreme noncoding conservation coincide with topologically associating domains (TADs) [9].

Numerous recent studies highlight and seek to elucidate the importance of functional non-coding regions, most recently by employing the CRISPR-Cas9 based techniques to locate and dissect elements that affect gene expression and phenotype/disease - associated processes [10–12]. Prioritizing target loci of interest for interrogating the function of their regulatory context will be one of the major focuses of functional genomic studies, as has been shown in the case of the *POU5F1* locus [13], and of *NF1, NF2* and *CUL3* genes [14]. CNEs and the regulatory landscapes defined by their clusters serve as excellent candidates for such studies [3,15,16].

A handful of CNE resources exist, mainly databases, which contain already pre-computed clusters of CNEs. These databases are static and mostly not updated. A summary of these resources is available in the review by Polychronopoulos et al. [6]. To our knowledge, there are only two tools available for the identification of conserved elements: PHAST [17] and CNEFinder [18]. The former relies on multiple sequence alignments and requires extensive computation time to derive “conserved” and “non-conserved” states from a two-state phylogenetic hidden Markov model (phylo-HMM), a space-time probabilistic model that considers both the nucleotide substitution of each base in the genome sequence through evolution and the transition from one base to the next. The latter produces CNEs based on a k-mer technique for computing maximal exact matches thus finding CNEs without the requirement of whole-genome alignments or indices. Neither of them comes with a comprehensive, easy-to-follow suite of tools tailored to the integrated exploration of CNEs from end-to-end: from identification to quality control and visualisation. Our package couples those processes together, enabling the user to harness the support and wealth of packages available through the common the Bioconductor infrastructure. Our package is specifically designed for efficient identification of CNEs using user-specified thresholds, and it functions equally well across vertebrates, invertebrates or plants. To study the evolutionary dynamics of these elements and their relationship to the genes around which they cluster, it is essential to be able to both produce and explore genome-wide sets of CNEs for a large number of species comparisons in a dedicated workflow, each with multiple length and conservation thresholds.

The *CNEr* package aims to detect CNEs and visualise them along the genome under a unified framework. For performance reasons, the implementation of CNE detection and corresponding I/O functions are primarily written as C extensions to R. We have used *CNEr* to produce sets of CNEs by scanning pairwise whole-genome net alignments with multiple reference species, each with two different window sizes and a range of minimum identity thresholds, available at http://ancora.genereg.net/downloads. In this work, we demonstrate the application of *CNEr* to the investigation of noncoding conservation between fruit fly *Drosophila* and tsetse fly *Glossina* - the two species at the evolutionary separation not previously investigated in insects [7] - and between two species of sea urchins. This has enabled us to observe some properties of GRB target genes shared across Metazoa. In a previous study, we showed that more distant comparisons in Diptera (between Drosophila and mosquitoes) failed to identify CNEs [7]. On the other hand, the conservation level across different species of the Drosophila genus is comparable to that across placental mammals. With *Drosophila* and *Glossina*, we wanted to explore the evolutionary divergence comparable to human vs. fish in another lineage and establish whether it is the same functional class of genes that is accompanied by CNEs featuring such a deep level of conservation. In the case of sea urchins, we wanted to investigate a lineage at an intermediate distance to vertebrates - closer than insects, more distant than the early branching chordates - in order to establish the continuum of GRBs across Metazoa. We present a series of downstream analysis of the newly identified CNEs, identifying their characteristic sequence features in invertebrates and functional classes of genes whose loci they span.

## Design and Implementation

### Overview of *CNEr* workflow

*CNEr* provides the functionality of large-scale identification and advanced visualisation of CNEs based on our previous strategies of detecting CNEs [7,8,19] as shown in Fig 1. *CNEr* scans the whole genome pairwise net alignment, which can be downloaded from UCSC or generated by the *CNEr* pipeline, for conserved elements. Various quality controls of the alignments are provided. The composition of aligned bases in the alignment can be used for tuning parameters during pairwise alignment (S1 Fig). More closely related species are expected to give higher rates of matched bases. The syntenic dotplot of the alignments (S2 Fig) quickly shows the syntenic regions between two assemblies.

**Fig 1:**
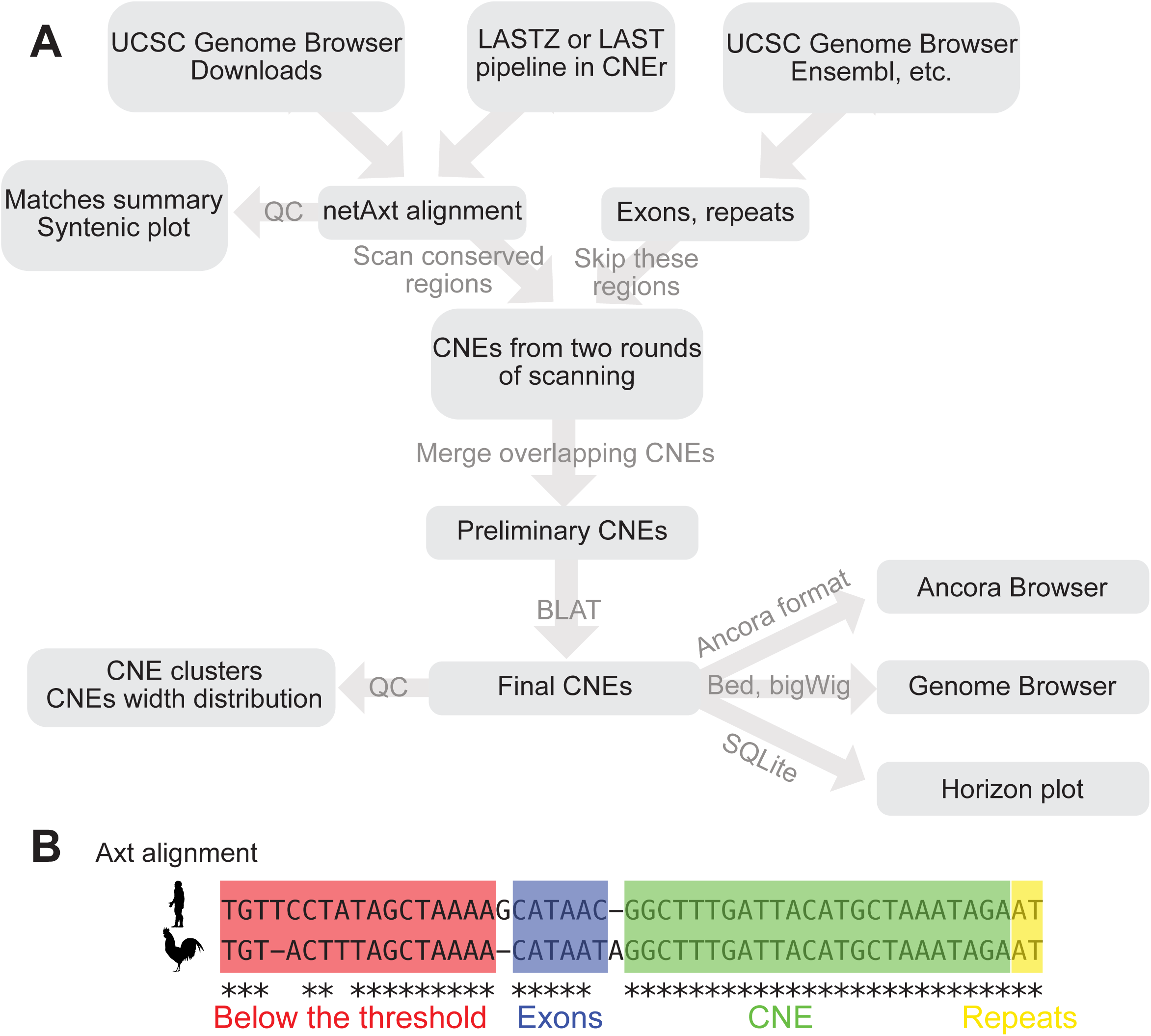
*CNEr* workflow. (A) A typical pipeline of identification and visualisation of CNEs. (B) Illustration of scanning an alignment for CNEs. The scanning window moves along the alignment for conserved regions. The exons and repeats regions are skipped during the scanning by default.

Considering the different extents of evolutionary divergence and sequence similarity between assemblies, we typically use the identity thresholds of 70% to 100% identity over a scanning window of 30 bp or 50 bp. Known annotations of exons and repeats are compiled from sources such as UCSC [20] and Ensembl [21] for common genomes, and elements overlapping with these regions are typically skipped during the scanning. Genome annotation pipeline, such as MAKER [22], can be used to create annotations for new genome assemblies.

Net alignments only keep the best match for each region in the reference genome. This is not acceptable when one of the aligned genomes underwent one or more whole genome duplications, leading to legitimate deviations from 1:1 orthology for many CNEs. To eliminate the bias of the choice of reference genome in the alignment and to capture duplicated CNEs during whole genome duplication (WGD), we scan two sets of net alignments by using each of the two compared genomes as reference in turn. This strategy performs well when comparing species with different numbers of WGD rounds, such as tetrapod vertebrates against teleost fish [23], or common carp [24] against other teleost fish. In such cases, some of the identified CNE pairs from two rounds of screening do overlap on both assemblies, and hence are merged into one CNE pair. As the last step, we align the CNEs back to the two respective genomes using BLAT and discard the ones with high number of hits. The remaining elements are considered to be a reliable set of CNEs.

*CNEr* provides a quick overview of the genomic distribution of CNEs along chromosomes. In S3 Fig, each CNE between human and mouse is plotted relative to each human chromosome (x-axis). We plot the cumulative number of CNEs over chromosomal positions. A CNE cluster is represented as a sharp increase of height in y-axis with small change in x-axis. For visualisation of CNEs in any genome browser, *CNEr* can export the CNE coordinates in BED file format and CNE density (measured by the percentage of area covered by CNEs within a smoothing window) in **bedGraph** and **bigWig** formats. Since running the whole pipeline of CNE detection can be time-consuming, we also implemented a storage and query system with SQLite as backend. Based on the visualisation capability of the *Gviz* package [25], *CNEr* can produce publication-quality horizon plots of CNE density along with other genomic annotations (see Methods and Data). Examples of the horizon plots are given in the following sections.

### *CNEr* package implementation

*CNEr* is a Bioconductor package developed in R statistical environment, distributed under the GPL-2 licence for *CNEr* code, and UCSC Kent’s licence for Jim Kent’s C source code it builds on [20]. Although *CNEr* supports compilation for both 32-bit and 64-bit systems across multiple platforms, it has limited functionality on the Windows platform due to the lack of the external sequence alignment software *BLAT* [26], which is required in the pipeline.

### Overview of whole genome pairwise alignment pipeline

UCSC Genome Informatics (http://hgdownload.soe.ucsc.edu/downloads.html) provides the pairwise alignments between many popular species. However, there is a frequent need to produce pairwise alignments for novel genome assemblies for new species, or using specific assembly versions when they are not available from UCSC. This pipeline mostly requires external sequence aligners and UCSC Kent’s utilities [20], and provides well-tested parameters for species with a varying degree of evolutionary divergence. In brief, first a sequence alignment software, *LASTZ* [27] or LAST [28], is used to find the similar regions between two repeat-masked genomes. Then, if two neighbouring alignments are close enough in the genome, they are joined into one fragment. During the alignment, every genomic fragment can match several others, and the longest one is kept. Finally, blocks of alignments are grouped into stretches of synteny and form the so called “net” alignments in Axt format [29]. *CNEr* comes with a vignette to demonstrate the whole pipeline. The produced Axt alignment can be manipulated in R as the *Axt* class, which is extended from *GRangePairs* class defined in *CNEr* (see S2 Text).

### Overview of the Axt scanning algorithm

The Axt alignment scanning algorithm constitutes the central part of this package for the identification of conserved noncoding elements. Due to the massive manipulation of characters, we implemented this algorithm purely in C for performance reasons; it is available to the R environment through R’s C interface. The minimal input is the Axt alignment and the ranges to filter out, i.e., the coding and/or repeat masked regions.

The Axt screening algorithm proceeds as in S1 Algorithm. First, the Axt alignment is converted into a linked Axt data structure as implemented in Jim Kent’s UCSC source code [20]. The filtering ranges are encoded into a hash table, where keys are the chromosome/sequences names and values are pointers to the linked lists of coordinates ranges. We then iterate over the linked Axt alignments. For each alignment, we use a running window to scan the alignment with a step size of 1 bp. Each base is searched against the filtering hash table and matched bases are skipped. All segments above the identity threshold are kept. The overlapping segments are merged into larger pieces. This procedure produces a set of CNEs conserved between the two aligned genome assemblies.

### *CNEr* visualisation capability

Instead of using the standard density plot for CNE density (as implemented in e.g. the Ancora browser), we introduce the horizon plot with the aim to increase the dynamic range of CNE density visualisation. The horizon plot provides a way of visualising CNE density over several orders of magnitude, and eliminates the need for multiple standard density tracks at different thresholds along the genomic coordinates. Instead, a relatively low conservation threshold is used, and multiple overlaid sections of the horizon plots will reveal peaks with different conservation density (see Fig 3A and Fig 3B in horizon plot, Fig 3C and Fig 3D in Ancora browser). We expand the functionality of “horizonplot” in *latticeExtra* package and integrate it into *Gviz* [25], which is the plot engine used in *CNEr*.

## Results

### *CNEr* use case I: *Drosophila-Glossina* CNEs

Here we demonstrate the application of *CNEr* to the analysis of Tsetse Fly (*Glossina morsitans*) CNEs and their putative target genes. *Glossina* is the sole vector of African trypanosomiasis (“sleeping sickness”), and it mediates transmission of the disease during feeding on blood. It has been shown previously [7] that, while there are tens of thousands of CNEs detected across different *Drosophila* species, there are almost no highly conserved elements found between *Drosophila* and malaria mosquito *Anopheles gambiae* or other mosquitos. *Glossina* and *Drosophila* are much closer to each other than either of them is to mosquitos, having a common ancestor that has diverged around 60.3 Mya (S4 Fig). With the recently available assembly and gene annotation of *Glossina* [30] (see S1 Text), we were able to identify clusters of CNEs between these two species. The clusters correspond to a subset of clusters defined by the CNEs derived from comparisons of different *Drosophila* species. A further investigation of gene functions, which are retained or missing in *Glossina*, was carried out by comparison with the *Drosophila* clusters.

A summary of CNEs detected between *Glossina* and *Drosophila* is given in Table 1. As expected, many fewer CNEs are detected from the comparison between *Glossina* and *Drosophila* than between any two *Drosophila* species, since *Glossina* is an outgroup to the *Drosophila*/*Sophophora* family. A closer examination of the CNE density plot in Ancora browser [31] revealed many missing clusters of CNEs relative to CNE density across *Drosophila* species, especially at a more stringent threshold. We wanted to find out if the missing and retained CNE clusters differ with respect to the functional categories of the genes they span. In the following analysis, the CNEs that are conserved for more than 70% over 30 bp are considered.

**Table 1:** The number of CNEs between D. melanogaster and several other species, including G. morsitans

| Minimum identity | vs *D.*<br>*ananassae* | vs *D.*<br>*pseudoobscura* | vs *D.*<br>*mojavensis* | vs *D.*<br>*virilis* | vs *G.*<br>*morsitans* |
| --- | --- | --- | --- | --- | --- |
| 70% over 30 bp | NC | NC | 176366 | 204970 | 9691 |
| 80% over 30 bp | NC | 313570 | 127293 | 146793 | 3924 |
| 90% over 30 bp | NC | 212951 | 81436 | 92288 | 1922 |
| 96% over 30 bp | 177759 | 128843 | 47408 | 52134 | 813 |
| 100% over 30 bp | 112073 | 76715 | 26972 | 29445 | 414 |
| 70% over 50 bp | 266385 | 248357 | 104476 | 120628 | 3185 |
| 80% over 50 bp | 223975 | 177266 | 66063 | 75204 | 1796 |
| 90% over 50 bp | 142899 | 96994 | 33455 | 37098 | 732 |
| 96% over 50 bp | 79631 | 49380 | 16387 | 17831 | 244 |
| 98% over 50 bp | 55460 | 33463 | 10741 | 11548 | 150 |
| 100% over 50 bp | 29218 | 17201 | 5250 | 5585 | 66 |
| NC, not counted due to the threshold being too low for close species. |  |  |  |  |  |

The most deeply conserved vertebrate CNEs are usually associated with genes involved in transcriptional regulation or development (trans-dev) functions [19]. Due to high divergence between *Drosophila* and *Glossina*, the regions with detectable CNE arrays tend to be of low CNE turnover, i.e. the process of sequence divergence and loss of ancestral CNEs is slow. If the same functional subset of genes is surrounded by low-turnover CNE clusters as in vertebrates, the encompassed genes will more likely be essential key developmental genes [5]. Indeed, *Drosophila* genes associated with (i.e. nearest to) *Glossina* vs. *Drosophila* CNEs are also associated with trans-dev terms (Fig 2A). Development, including organ, system and tissue development, appears at the majority of the top Gene Ontology (GO) terms. The other highly significant GO terms include biological regulation, regulation of cellular process and cell differentiation. CNE clusters can span regions of tens to hundreds of kilobases around the actual target gene, which is on average shorter than the equivalent spans in vertebrate genomes. This is in agreement with our observation that CNE clusters and the GRBs they define (and, by extension, the underlying TADs) expand and shrink roughly in proportion to genome size [9]. The *H15* and *mid* locus (Fig 3A) is one of the biggest CNE clusters retained between *Glossina* and *Drosophila*. The *H15* and *mid* genes encode the T-box family proteins involved in heart development [32]. Although the CNE density between *Drosophila* and *Glossina* is much lower than that across the *Drosophila* genus, it clearly marks the CNE cluster boundaries of this locus, containing 67 CNEs at the 70% identity over 30 bp threshold. For the 40 largest retained CNE clusters, we provide a comprehensive list of CNE cluster coordinates, the target genes, the protein domains and the number of associated CNEs (S1 Table). As we can see, the majority of the target genes have *Homeobox, Forkhead* or *C2H2* Zn finger domains, just like the genes spanned by the most conserved CNE clusters in vertebrates.

**Fig 2:**
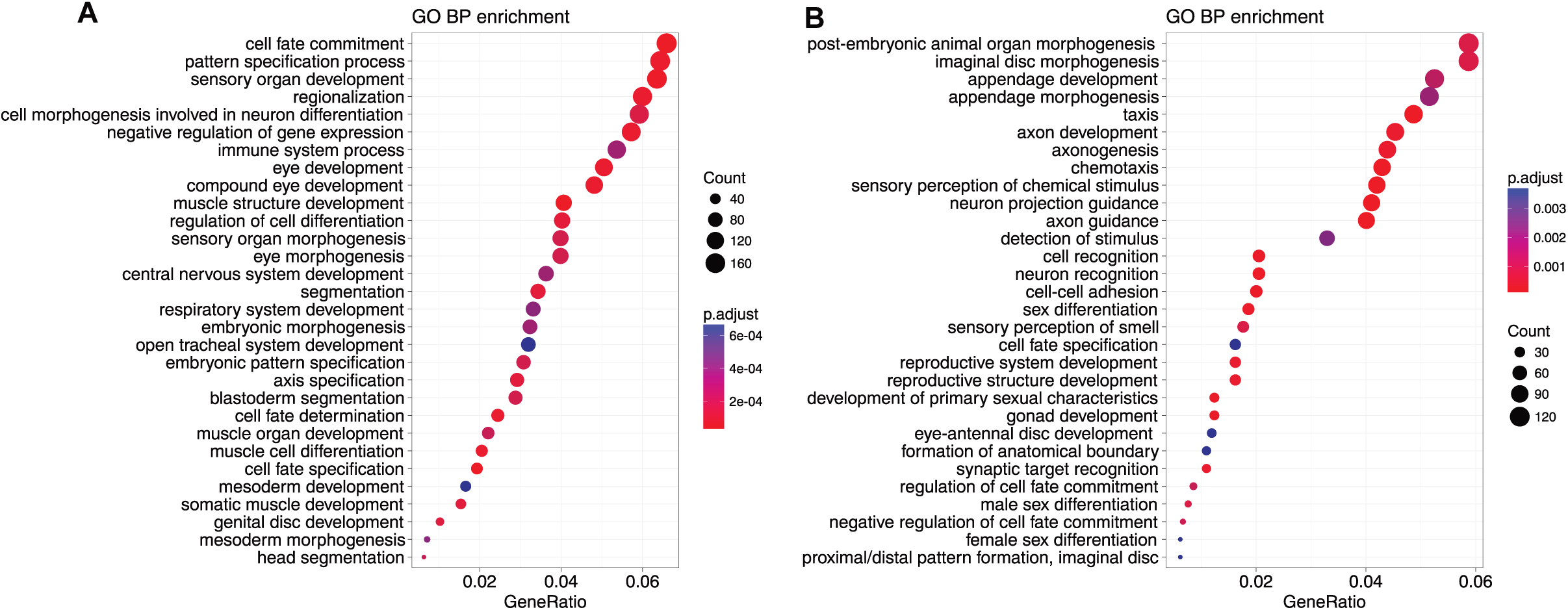
Over-represented GO Biological Process terms ranked by GeneRatio. The gene ratio is defined as the number of genes associated with the term in our selected genes divided by the number number of selected genes. The p-values are adjusted using “BH” (Benjamini-Hochberg) correction. The visualisation is done by *clusterProfiler [33]*. (A) GO enrichment for genes nearest to *Drosophila* and *Glossina* CNEs. (B) GO enrichment for genes in the missing CNEs clusters compared between *Drosophila* and *Glossina*.

**Fig 3:**
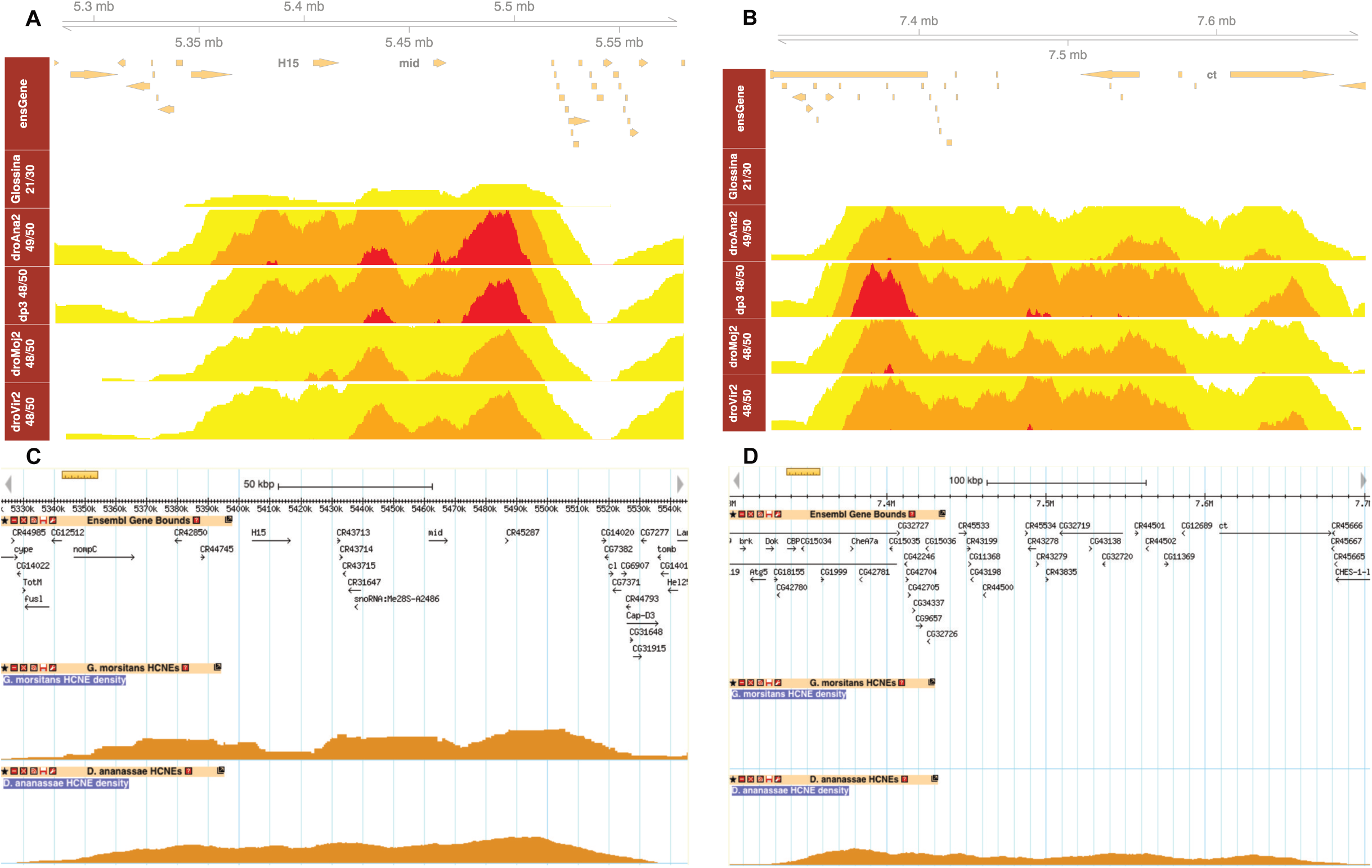
Horizon plot of CNE density around key developmental genes along *D. melanogaster* as reference. (A) *H15* and *mid* genes are spanned by arrays of CNEs. Despite the much lower CNE density from *D. melanogaster* and Glossina, a CNE cluster boundary shows up that is consistent with CNEs from other drosophila species. (B) The CNE cluster around *ct* gene is missing in the comparison of *D. melanogaster* and Glossina since no CNEs are detected. This implies that this region undergoes a higher CNE turnover rate. (C, D) The same loci as in (A, B) are shown on the Ancora browser in order to compare the normal CNE density plot with the horizon plot. Notations: ensGene, Ensembl gene track; *Glossina* 21/30, G. morsitans 70% identity over 30 bp; droAna2 49/50, *D. ananassae* 98% identity over 50 bp; dp3 48/50, *D. pseudoobscura* 96% identity over 50 bp; droMoj2 48/50, *D. mojavensis* 96 % identity over 50 bp; droVir2 48/50, *D. virilis* 96% identity over 50 bp.

Some other regions have strong clusters of CNEs conserved among *Drosophila* species, however, the CNE cluster between *Drosophila* and *Glossina* is absent. The *ct* locus (Fig 3B), encoding the cut transcription factor, is one of the best representative cases. Ct plays roles in the later stages of development, controlling axon guidance and branching in the development of the nervous system, as well as in the specification of several organ structures such as Malpighian tubules [34]. In order to locate the CNE clusters missing from *Drosophila* vs. *Glossina* comparison, we use the CNE clusters detected in *D. melanogaster* vs. *D. ananassae* comparison as reference and compare them with the aforementioned retained CNE clusters. The genes within those missing CNE clusters are highly enriched for axon guidance and neuronal development (Fig 2B). We then examine the CNE turnover rate (the speed of replacing old CNEs) of the 216 human genes that are associated with the axon guidance term (GO:0007411), with both human and *Drosophila* as reference. The turnover rate is calculated as the reduction of the number of CNEs between two sets of CNEs. For human reference, we choose the CNE set of human vs. mouse and human vs. zebrafish, while *D. melanogaster* vs. *D. ananassae* and *D. melanogaster* vs. *Glossina* are chosen for *Drosophila* reference. As shown in Fig 4, the axon guidance genes have significantly higher turnover rate than the other genes (p < 1e-5, Kolmogorov-Smirnov one-sided test) in both human and *Drosophila* lineages.

**Fig 4:**
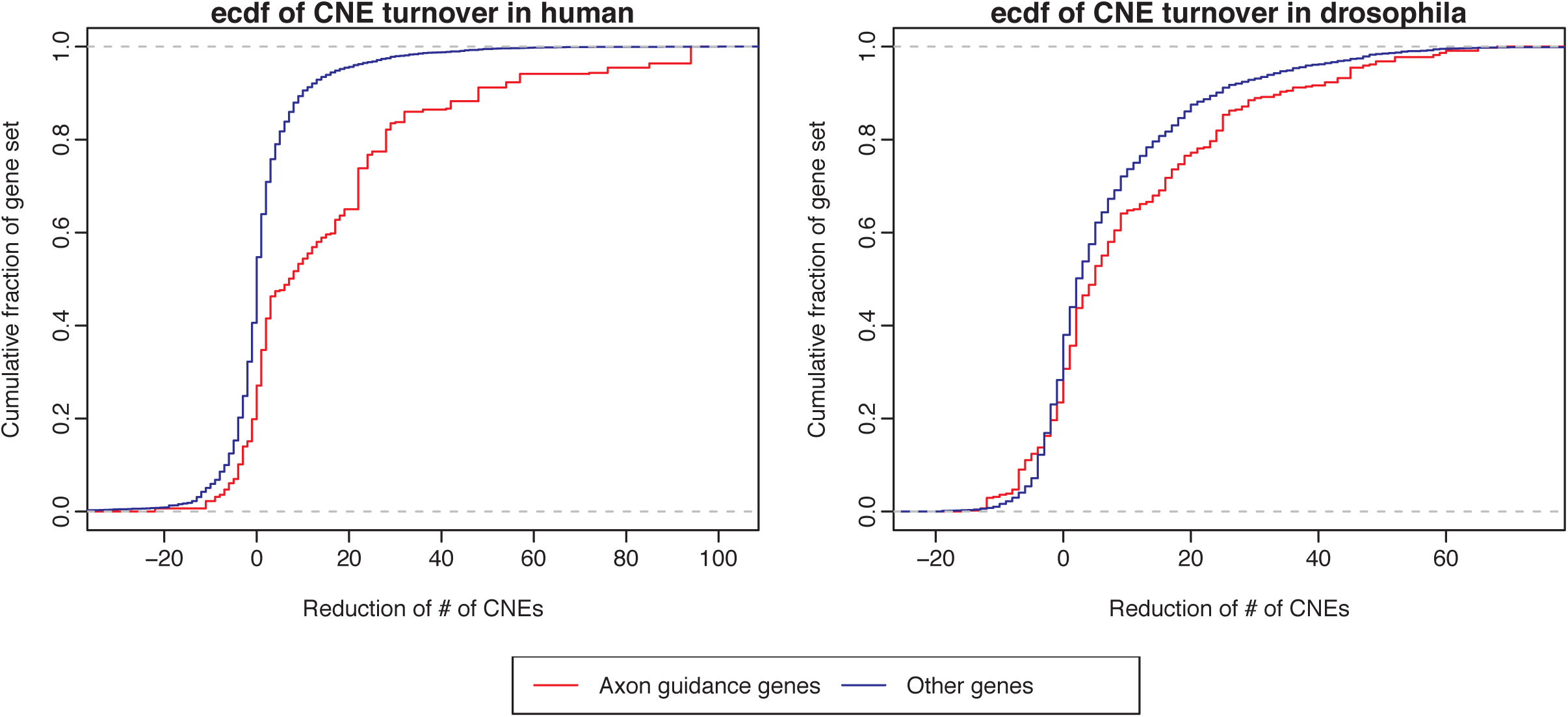
Cumulative distribution function of the changes of CNE number. For a 40kb window around each orthologous gene pair between human and *drosophila*, we calculate the reduction of the number of CNEs for human (# of CNEs from human-mouse comparison minus # of CNEs from human-zebrafish comparison) and drosophila (# of CNEs from *D. melanogaster* vs. *D. ananassae* comparison minus # of CNEs from *D. melanogaster* vs. *Glossina*) as reference. The axon guidance genes significantly show a higher degree of CNE number reduction, compared with the other genes (p < 1e-5, Kolmogorov-Smirnov one-sided test).

### *CNEr* use case II: sea urchin CNEs

In this section we apply *CNEr* to the comparison of highly fragmented genome assemblies of two sea urchin species *Strongylocentrotus purpuratus* and *Lytechinus variegatus* (see S1 Text). The purpose of this analysis is twofold. First, we want to demonstrate how well *CNEr* is able to call CNEs and their clusters in the case of highly fragmented draft genomes: the ability to perform this analysis on draft genome assemblies would show that our approach can be applied to a large number of available genomes, most of which haven’t been assembled past the draft stage and are likely to remain in that state. Second, we wanted to ask if a third lineage, evolutionarily closer to vertebrates than insects but still lacking any shared CNEs with vertebrates, would exhibit the same patterns of noncoding conservation. This could provide a hint towards CNEs’ universal presence in Metazoa, in addition to providing an informative additional dataset for comparative studies of genomic regulatory blocks.

*S. purpuratus* is a popular model organism in cell and developmental biology. *S. purpuratus* and *L. variegatus* have a divergence time of 50 Mya [35] and historically moderate rates of sequence divergence, which makes them ideal for comparative genomics studies of regulatory elements. We identified 18,025 CNEs with threshold of 100% identity over 50 bp window. Despite the highly fragmented assemblies, we could clearly detect 808 prominent CNE clusters. An especially interesting observation is the largest cluster we detected, at the *Meis* gene locus (Fig 5). The CNE density clearly marks the boundaries of the CNE cluster. In Metazoa, *Meis*, one of the most well-known homeobox genes, is involved in normal development and cell differentiation. Tetrapod vertebrates have three *Meis* orthologs as a result of two rounds of whole genome duplication. The CNE cluster around *Meis2* (one of three Meis paralogs arisen by two WGD rounds at the root of vertebrates) is the largest such cluster in vertebrates [19]. Remarkably, the cluster of CNEs around *Drosophila*’s *Meis* ortholog, *hth* (homothorax), is also the largest CNE cluster in the *D. melanogaster* genome [7]. It is currently unknown why the largest clusters of deeply conserved CNEs are found around the same gene in three different metazoan lineages, even though none of the CNEs from one lineage has any sequence similarities to CNEs in the other two. The most plausible explanation is that the ancestral *Meis* (*hth*) locus was already the largest such locus in the ancestral genome, and that CNE turnover has led to three separate current lineage-specific sets of CNEs.

**Fig 5:**
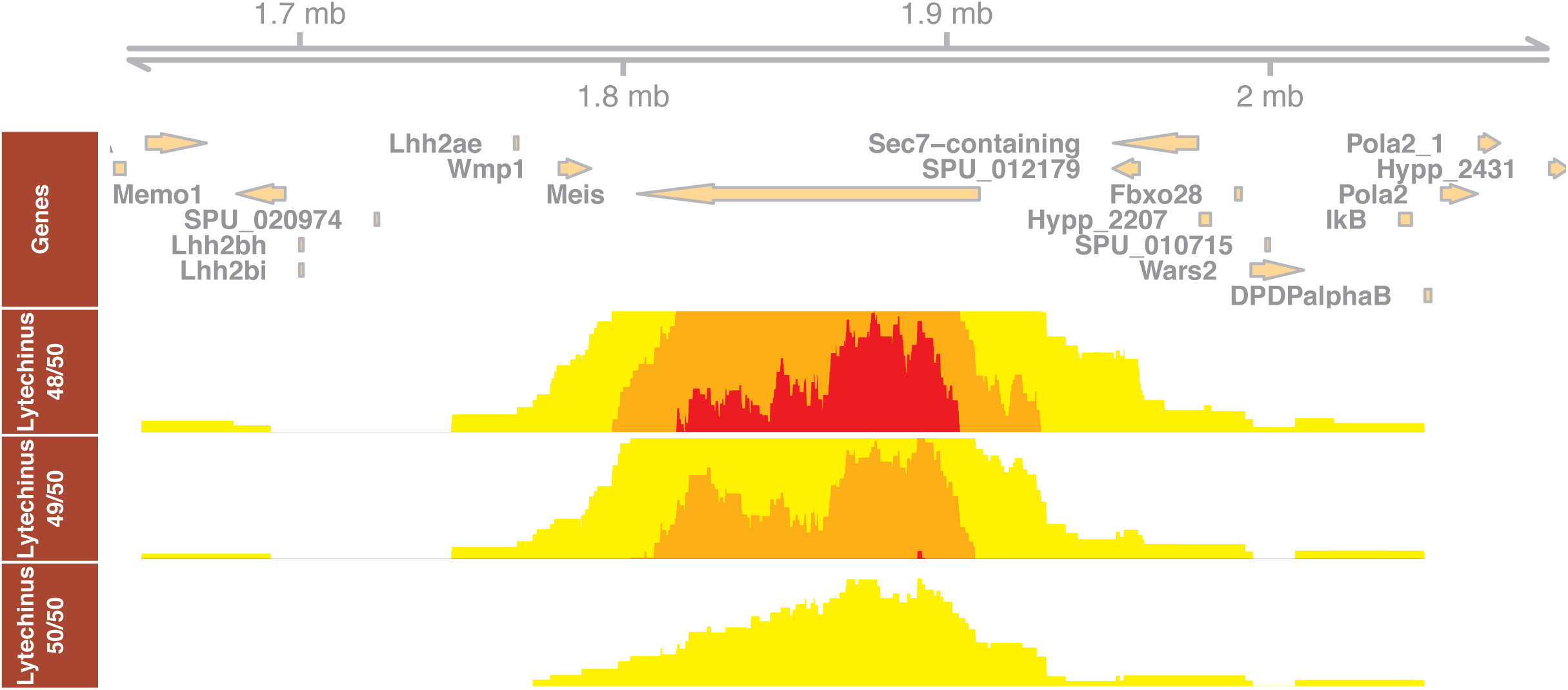
Horizon plot of CNE density at *Meis* loci on sea urchin *Strongylocentrotus purpuratus.* The density plots of CNEs detected at similarity threshold 96% (48/50), 98% (49/50) and 100% (50/50) over 50 bp sliding window, in *Lytechinus variegatus* comparison, are shown in three horizon plot tracks. The boundaries of CNE clusters from various thresholds are mutually consistent.

### *CNEr* reveals interesting sequence features characteristic of ultraconservation

It has been shown that vertebrate nonexonic CNEs are enriched in the TAATTA hexanucleotide motif, which is an extended recognition site for the homeodomain DNA-binding module [36]. With *CNEr*, we can easily verify the existence of TAATTA motif in CNEs of invertebrate species. In S5 Fig A, we consider CNEs identified by *CNEr* that are conserved between *D. melanogaster* and *D. virilis* over 98% for more than 50 nucleotides and plot them by increasing width using *heatmaps* package (https://bioconductor.org/packages/heatmaps/). The first two heatmaps confirm that CNEs are enriched in AT inside but exhibit a marked depletion of AT at their borders, consistent with what is known about their biology in vertebrates [37]. Furthermore, the TAATTA motif is enriched in insect CNEs. The motif seems to be extended further by flanking A/T nucleotides. When replacing A/T (W) with G/C (S), the heatmap pattern disappears. We ask whether this is a general property of CNEs in Metazoa and, using *CNEr*, proceed to the identification of CNEs that are conserved between (a) *C. elegans* and *C. briggsae* at 100% for more than 30 nucleotides (worm CNEs, see S5 Fig B), (b) *L. variegatus* and *S. purpuratus* at 100% for more than 50 nucleotides (sea urchin CNEs, see S5 Fig C). We observe that the same pattern does not hold in those cases, i.e. it appears like enrichment of CNEs in TAATTA is not a universal phenomenon but applies only to insect and vertebrate elements. Our pipeline is a powerful tool for studying the question of how and when this TAATTA-enrichments originated, as well as a multitude of related questions.

### Downstream overlap analysis of CNEs reveals that they are highly enriched in stem-cell regulatory elements

Finally, we demonstrate the utility of CNEr for general hypothesis generation about CNEs by identifying elements highly conserved (>98% identity over 50 bp) between human and chicken and performing global and pairwise overlap analyses against various genomic features using the R/Bioconductor packages *LOLA* [38] and *regioneR* [39], respectively. *LOLA* allows for enrichment analysis of genomic intervals using a core reference database assembled from various resources of genomic data, while *regioneR* permits cross-validation of the findings through pairwise overlap analyses. As evident from inspection of S2 Table and Fig 6, both packages converge to the conclusion that the identified CNEs between human and chicken are significantly enriched in *Sox2* and *Oct-4* (*POU5F1*) binding sites. *Sox2* and *Oct-4*, in concert with Nanog, are believed to play key roles in maintaining pluripotency. This finding comes in accordance with previous reports suggesting that several CNEs are enriched in classical octamer motifs recognized by developmental homeobox transcription factors [40]. Nonetheless, this is the first time that such an association of the most deeply conserved CNEs with key pluripotency elements is reported in the literature, and we anticipate that more associations of this kind will be revealed in the future by coupling *CNEr* with other R/Bioconductor packages.

**Fig 6:**
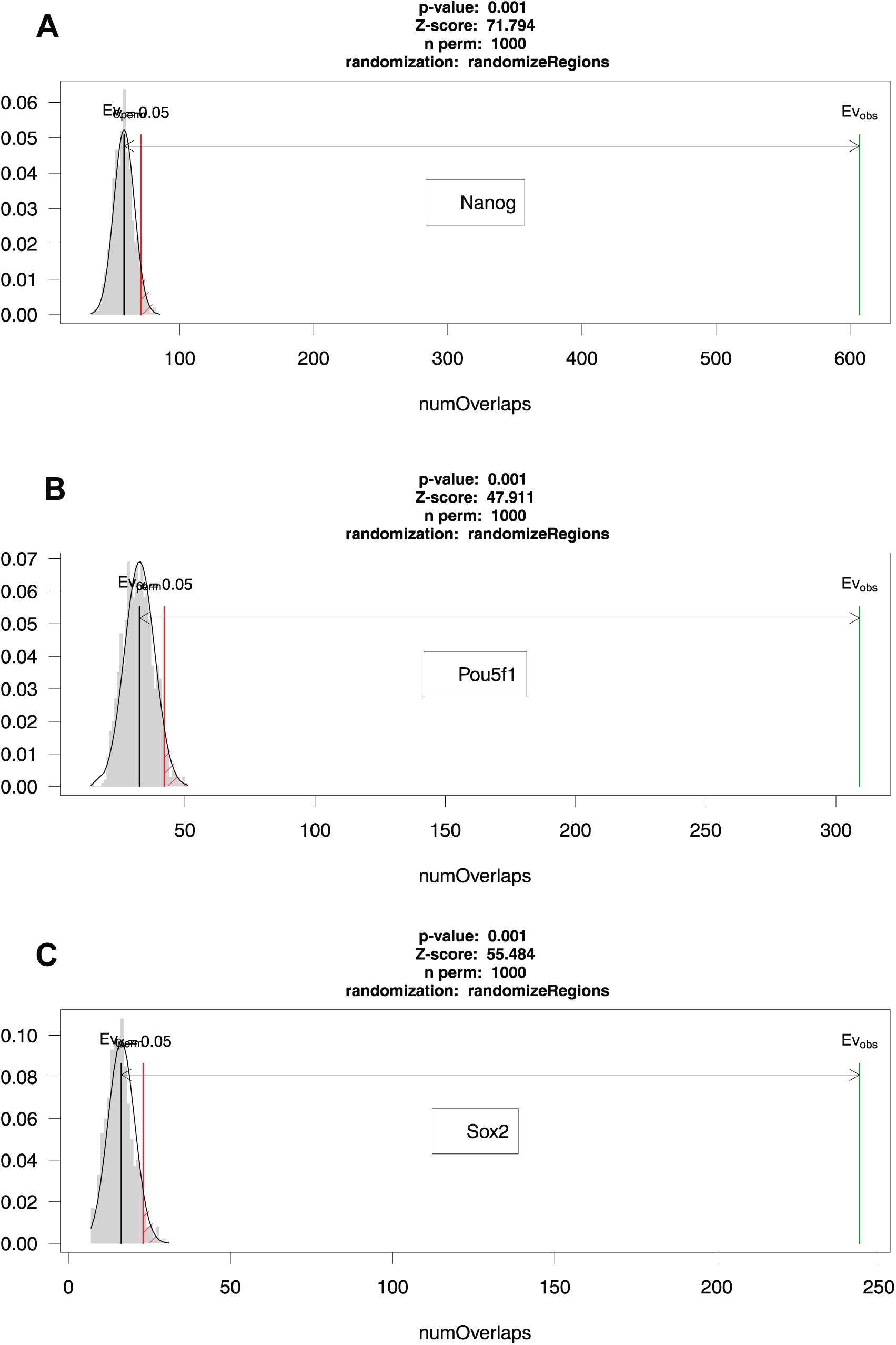
Pairwise overlap analysis of CNEs demonstrates association with *Sox2, POU5F1* and Nanog binding regions. In all three cases, permutation tests with 1000 permutations of CNEs are shown. In grey the number of overlaps of the randomized regions with the test regions of interest (in this case, *Sox2, POU5F1* and *Nanog*) are depicted. Those overlaps of the randomized regions cluster around the black bar that represents the mean. In green, the number of overlaps of the actual regions (CNEs in this case) with the test regions is shown and is proved to be much larger than expected in all cases. The red line denotes the significance limit.

### Availability and Future Directions

The *CNEr* package with self-contained UCSC Kent’s utility source code is available at Bioconductor release branch http://bioconductor.org/packages/CNEr/. Active development and bug reports is hosted on github https://github.com/ge11232002/CNEr/. Currently the *BLAT* is the only supported aligner for identifying repeats. Other high performance short read aligners that run in parallel, such as Bowtie1/2 and BWA, are desired for large set of CNEs. Furthermore, integration of GRB identification approach and GRB target gene prediction is planned for future development.

## Declarations

### Authors’ contributions

GT and BL conceived the project. GT implemented the software, analyzed the data. DP tested the software, analyzed data. GT, DP, BL wrote the manuscript.

#### Acknowledgements

We are grateful to the Bioconductor community for trying out the *CNEr* package and providing useful input.

## Supporting information

S1 Text. Glossina and sea urchin data.

S2 Text. Working with Paired Genomic Ranges

S1 Fig. The heatmap shows the percentage of matched bases in the Axt alignments.

This can be useful for examining the quality of Axt alignments, especially from the whole genome pairwise alignment pipeline in *CNEr* package. The left panel has higher rates of matches than right panel since the divergence of human and mouse is much smaller than that between human and zebrafish.

S2 Fig. The syntenic plot of alignment blocks between chr1, chr2 of human and chr1, chr2 of mouse.

This plot is mostly used for tuning the parameters during whole genome pairwise alignment to get better alignments. It can also show ancient duplications for the alignment of a sequence against itself.

S3 Fig. The distribution of CNEs along the 6 biggest chromosomes in human genome.

Each CNE is plotted as a dot with the position in chromosome as x-axis. A sharp increase in y-axis represents a CNE cluster.

S4 Fig. The species tree of Drosophila, Glossina and mosquitos.

The phylogenetic tree is constructed based on the data on last common ancestors from TimeTree (Hedges et al., 2006). The genome of the malaria mosquito *A. gambiae* is highly divergent from *Drosophila* family and unsuitable for comparative genomics study, while *G. morsitans* is much closer.

S5 Fig. Sequence heatmaps of CNEs in different lineages.

(A) *D. melanogaster* and *D. virilis* (B) *C. elegans* and *C. briggsae* (C) *L. variegatus* and *S. purpuratus.* The CNEs are ranked by decreasing CNE width.

S1 Table. A list of the most prominent CNE clusters detected between *Drosophila* and *Glossina*.

S2 Table. Overlap analysis of CNEs.

Global overlap analysis of CNEs against multiple genomic features using LOLA reveals that they overlap with Nanog, Sox2 and POU5F1 binding regions. This table is the output of runLOLA algorithm sorted by FDR. All top hits include important factors associated with pluripotency. userSet: all CNEs conserved between human and chicken over 98% identity for more than 50 bp. collection: all elements in CODEX database. universe: all active DNase I hypersensitive sites.

S1 Algorithm. The algorithm of scanning Axt alignment.

